# Cell segmentation and gene imputation for imaging-based spatial transcriptomics

**DOI:** 10.1101/2023.10.26.564185

**Authors:** Yunshan Zhong, Xianwen Ren

## Abstract

Imaging-based spatial transcriptomics technologies are revolutionary tools for biomedical investigation, but the power is currently limited by small number of measured genes and tricky cell segmentation. Here we introduce RedeFISH to simultaneously conduct cell segmentation and gene imputation for imaging-based spatial transcriptomics with the aid of single-cell RNA sequencing data. Extensive benchmarking across various spatial platforms and tissue types shows the validity and power of the cell-segmented, whole-transcriptome spatial data generated by RedeFISH.

## Main text

The rapid development of spatial transcriptomics (ST) technologies has provided unprecedented resolution to investigate spatial gene expression. In addition to ST platforms, *e*.*g*., 10x Genomics Visium, which measure gene expression at spatial resolution of multiple micrometers^1, 2^, single-molecule ST platforms achieve subcellular resolution by localizing mRNAs in situ^3-5^. Such higher resolution and limited number of genes detected raise new analytical questions, with cell segmentation and gene imputation typically as the foremost post-experiment steps to obtain the cellular and whole-transcriptome characteristics pivotal to systematic investigation.

Although multiple methods have been developed for cell segmentation, these methods depend on morphologically-defined cells with additional staining images^6-9^. Because such algorithmic configuration requests additional data including nuclei-or membrane-staining images and manual cell typing for training and learning, the applications are technically challenging, labor-intensive, and time-consuming, limiting the broad application of ST technologies. In particular, the rapid development of ST technologies can now generate ultra-huge datasets that comprise billions of transcripts for centimeter-scale tissues, raising the necessity of automatic and efficient analytical tools. In addition, the small number of genes measured by imaging-based ST technologies also greatly limits biological discoveries.

As single-cell RNA sequencing (scRNA-seq) has been widely applied and scRNA-seq data contain inherently the information of cellular identity and whole-transcriptome expression, here we propose an algorithm named as RedeFISH to automatically conduct cell segmentation and gene imputation for imaging-based ST data with the aid of scRNA-seq data.

Different from the previous “morphologically-defined cell segmentation” algorithms, RedeFISH implements “functionally-defined cell segmentation” by maximizing the alignment of ST and scRNA-seq data at the cellular resolution.

The functional consistency between ST segments and scRNA-seq cells resulted from RedeFISH enables transfers of whole-transcriptome expression profiles of scRNA-seq data to precise locations of ST data, generating cell-segmented, whole-transcriptome spatial data that are ideal for biological investigation.

In brief, we define a Segmentation Module to generate segments from ST data and a Deconvolution Module to identify the optimal scRNA-seq cells matched with those ST segments (Fig.1a). With iteration between the two modules within a reinforcement learning framework, RedeFISH optimizes ST segmentation by maximizing the matching degrees between ST segments and scRNA-seq cells (Fig. 1a), without the need of additional staining data, cell-type labels, or training.

**Fig 1.**
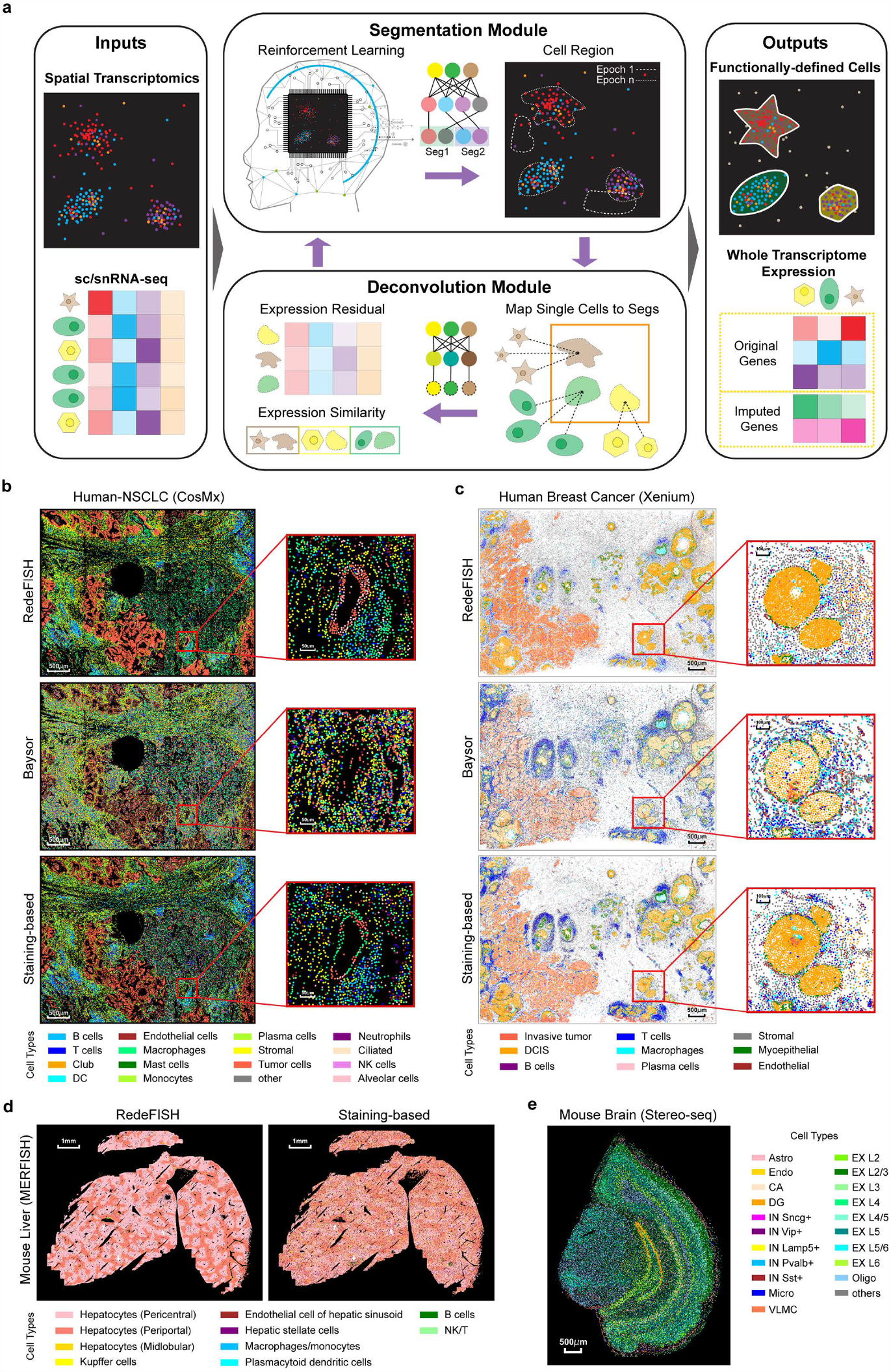
Overview of the RedeFISH algorithm and spatial distribution of cell types. **a**, Overview of RedeFISH workflow. RedeFISH requires sc/snRNA-seq data together with spatial transcriptomics data as input and adopts an iterative approach for optimizing functionally-defined cell regions (Segmentation Module) and aligning single cells and these regions (Deconvolution Module), with feedbacks providing to each other. The final output of the model consists aligned cell regions and their corresponding whole-transcriptome expression profiles. **b**, Spatial distribution patterns of cell types within the Human-NSCLC (CoxMx) dataset through label transferring of identified cells from RedeFISH, Baysor and Staining-based method. DC, dendritic cell; NK, natural killer. **c**, Spatial distribution patterns of cell types within the Human Breast Cancer (Xenium) dataset through label transferring. DCIS, ductal carcinoma in situ. **d**, Spatial distribution patterns of cell types within the Mouse Liver (MERFISH) dataset through label transferring. NK, natural killer. **e**, Spatial distribution patterns of cell types within the Mouse Brain (Stereo-seq) dataset through label transferring. Astro, astrocyte; CA, cornu ammonis area; DG, dentate gyrus; Endo, endothelial cell; EX, excitatory glutamatergic neuron; IN, GABAergic interneuron; Micro, microglia; Oligo, oligodendrocyte; VLMC, vascular and leptomeningeal cells.

We evaluated the performance of RedeFISH on six ST datasets across four representative platforms including MERFISH, Xenium, CosMX, and Stereo-seq, different species including human and mouse, and different organs including ileum, breast, brain, liver and lung. Compared with the morphologically-defined cell segmentation results from Baysor, a state-of-the-art (SOTA) segmentation algorithm^10^, and the manual segmentation results published in the original papers with the help of staining data, RedeFISH exhibited stable superior performance regarding the quality of identified cells and the scalability to handle ultra-huge ST datasets. These results prove the validity of functionally-defined cell segmentation, and suggest the feasibility to obtain cell-segmented, whole-transcriptome spatial data by automatic cell segmentation and gene imputation of imaging-based ST data with the aid of scRNA-seq.

First, RedeFISH achieves excellent alignment between ST segments and single cells in scRNA-seq data regarding gene expression similarity. RedeFISH exhibited superior performance compared to both Baysor and staining-based methods, with significantly higher cosine similarity for identified cells across all six evaluated datasets (Extended Data Fig. 1) and significantly lower root mean squared error (RMSE) values (Extended Data Fig. 2) obtained. Dividing ST 2D space grids and assigning matched cells to the nested grids further confirmed the superperformance of RedeFISH, demonstrating the high quality of functionally-defined cells (Extended Data Fig. 1 and Extended Data Fig. 2).

RedeFISH demonstrates great computational efficiency compared with previous algorithms (Extended Data Figure 3).

Across a wide spectrum of dataset sizes, ranging from ~1 million to ~100 million transcripts, RedeFISH exhibited approximately 10-fold improvement than Baysor. In case of ultra-large datasets (>100 million transcripts), RedeFISH accomplished cell segmentation and gene imputation within one day. However, alternative approaches, such as Baysor, failed for such tasks due to computational limitations.

We next illustrate how RedeFISH reveals spatial structures through cell segmentation of ST data with the aid of scRNA-seq (Fig.1b-e and Extended Data Fig. 4-9). We first evaluated RedeFISH on a human non-small cell lung cancer (NSCLC) dataset generated by the Nanostring CosMx platform^4^, which captured ~150 mm^2^ of lung tissue with identifiable tumor regions and tertiary lymphatic structures. RedeFISH successfully revealed the alveoli structures, of which tumor cells and alveolar cells formed circles (Fig. 1b and Extended Data Fig. 4). Staining-based methods also revealed alveoli-like circular structures, but Baysor and staining-based methods failed to reveal intact alveoli at certain position as we exemplified in Fig.1b. Close to the alveoli we highlighted, RedeFISH and staining-based methods, but not Baysor, revealed a region enriched by T and B cells, which was potentially equivalent to the tertiary lymphatic structure defined pathologically (Fig. 1b and Extended Data Fig. 4). Because of the whole-transcriptome feature of scRNA-seq data, RedeFISH also imputed the spatial distributions for genes that did not have probes designed by the CosMx platform (Extended Data Fig. 4), further confirming the nature of the spatial structures revealed by RedeFISH.

Then we evaluated the performance of RedeFISH on a human breast cancer dataset generated by the 10x Genomics Xenium platform^5^, which comprises a HER2+/ER+/PR− tissue block and contains multiple regions of ductal carcinoma in situ (DCIS) and invasive tumors. RedeFISH, Baysor, and staining-based manual curation all identified circular duct-like structures, with DCIS cells crowded to each other and surrounded by myoepithelial cells (Fig.1c and Extended Data Fig. 5), confirming the effectiveness of functionally-defined cell segmentation on the Xenium platform. Different from RedeFISH, staining-based curation identified more unexpected aberrant “cells” characterized by both extremely low density of Xenium transcripts and irregular nuclei morphology localized at the middle portion of the duct we highlighted (Extended Data Fig. 5b). Baysor, otherwise, reported more cell types beyond the DCIS cells in the basal region of the duct without histological or transcriptomic supports (Extended Data Fig. 5b).

On a mouse liver dataset generated by the MERFISH platform^10^, of which a total of 347 genes were profiled, RedeFISH also yielded excellent performance similar to that of staining-based algorithms, with boundaries between the pericentral and periportal regions of hepatocytes clearer in RedeFISH segmentation (Fig 1d and Extended Data Fig. 6).

Similarly, RedeFISH can also be applied to ST data generated by the Stereo-seq platform. On a mouse brain dataset generated by Stereo-seq^11^, with cortex and subcortical regions of various neuronal and non-neuronal cells included, RedeFISH successfully revealed the hierarchy pattern of different cell types, including excitatory glutamatergic neurons and several non-neuronal cells (astrocytes, oligodendrocytes, microglia etc) (Fig.1e and Extended Data Fig. 7).

We further illustrated the performance of RedeFISH with H&E images as references. Four regions of interest (ROIs) across different tissue regions of the Human Breast Cancer (Xenium) dataset were zoomed in for comparison (Extended Data Fig. 10a). Macrophages were predicted to occupy nearly the entire duct in ROI #1 by RedeFISH (Extended Data Fig. 10b), consistent with the H&E images, while Baysor generated more fragments in contrast. In ROI #2, RedeFISH successfully identified the circular vascular structures surrounded by endothelial cells (Extended Data Fig. 10b), consistent with H&E images, while Baysor and the staining-based method failed to identify such structures. Similarly, RedeFISH successfully identified vascular structures in ROI #3 and invasive tumor cells in ROI #4, while Baysor failed (Extended Data Fig. 10b).

Cell segmentation of ST data with the aid of scRNA-seq by RedeFISH not only enables automatic functionally-defined cell segmentation, but also enables whole-transcriptome spatial expression analysis that is impossible based on ST data only. Wnt/β-catenin signaling has been shown to play important roles in intestinal stem cell maintenance, differentiation and upon deregulation^12^, but *in situ* investigation of Wnt/β-catenin signaling directly based on ST data is currently difficult. For example, a mouse ileum ST dataset generated by the MERFISH platform profiled the spatial distribution of 241 genes, of which 12 genes were related to Wnt/β-catenin signaling (11.3% covered, according to the Gene Ontology annotation). We applied RedeFISH to this mouse Ileum dataset, identified the clear histological structures consistent with intestinal anatomy via functionally-defined cell segmentation (Fig.2a-b). The ST segments generated by RedeFISH showed consistent marker gene expression similar to scRNA-seq observation (Extended Data Fig. 11a). In addition to cell markers explicitly designed by MERFISH, RedeFISH automatically upgraded the spatial data of 241 genes to the whole transcriptome, and identified more genes that are cell-type-specific and spatially informative (Fig. 2a). With the imputation by RedeFISH, *in situ* investigation of the spatial regulatory mechanisms of Wnt/β-catenin signaling in intestinal stem cell maintenance and differentiation becomes possible (Extended Data Fig. 11b). Consistent with previous studies, *Wnt3* (not measured by MERFISH but imputed by RedeFISH) was highly expressed in Paneth cells^13^ (Fig.2c-d). *Fzd5*, encoding a receptor of *Wnt* ligand and measured by MERFISH, was highly expressed in stem and progenitor cells (Fig.2c-d). The spatial distribution of *Wnt3* (imputed by RedeFISH) and *Fzd5* (measured by MERFISH) showed the potential regulatory roles of Paneth cells in intestinal stem cell maintenance and differentiation, consistent with previous finding that Paneth cells are essential components of intestinal stem cell niches^14^. The whole-transcriptome imputation by RedeFISH enables *in situ* investigation of more spatial regulatory mechanisms, as exemplified by the diverse combinatory interactions between *Wnt*-genes-encoding molecular components and and their ligands/receptors (Extended Data Fig. 11b).

**Fig 2.**
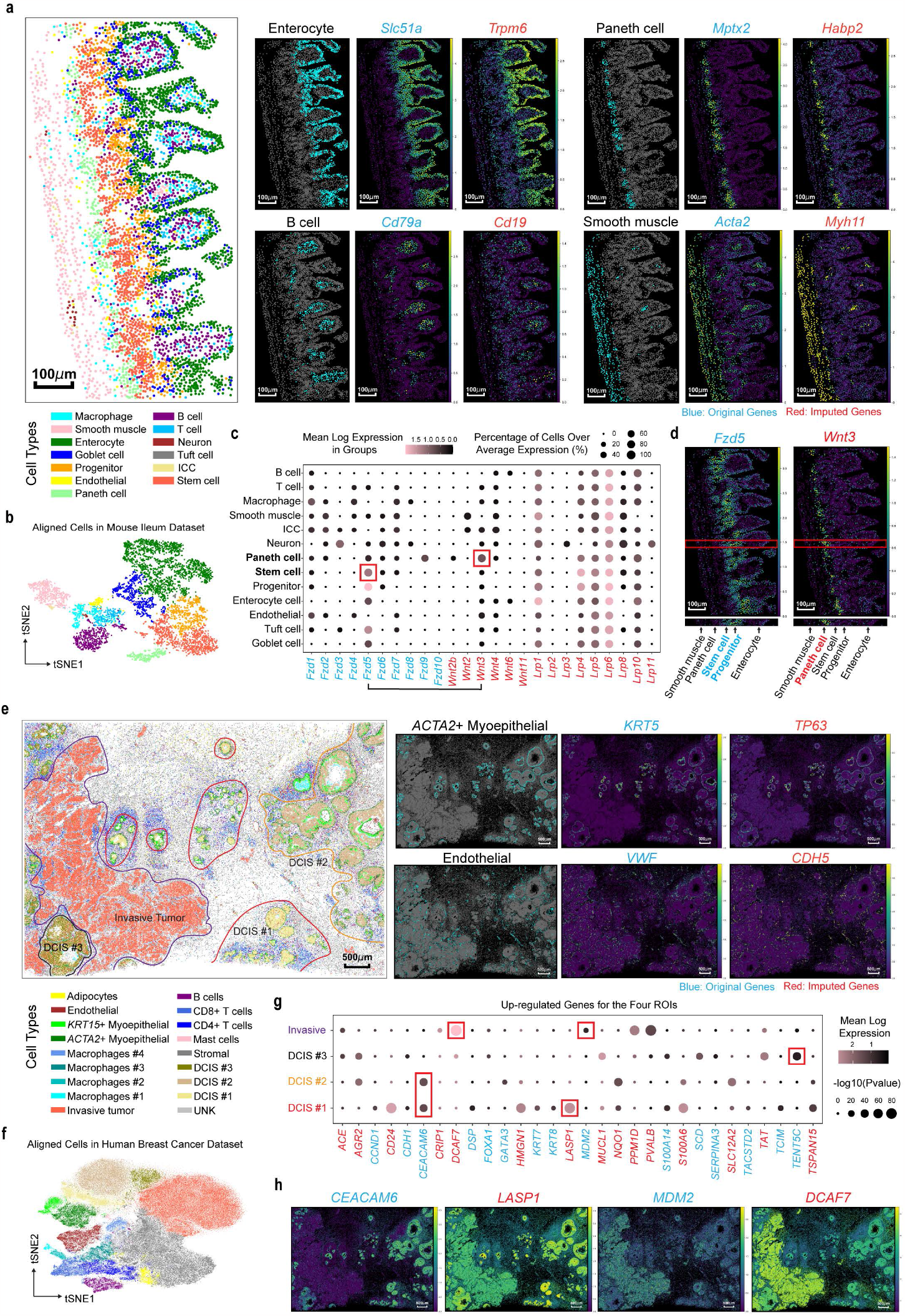
RedeFISH enables analysis of intercellular communication and tumor microenvironment. **a**, Spatial distribution pattern of cell types and marker genes within the Mouse Ileum (MERFISH) dataset. Left: Cell-type distribution through manual annotation. Right: Distribution of enterocytes, Paneth cells, smooth muscle cells, and B cells, along with their corresponding original and imputed marker genes. ICC, Interstitial cells of Cajal. **b**, tSNE plot of the cell types in Mouse Ileum dataset. **c**, Expression of *Wnt, Fzd* and *Lrp* in cell types. The size of the dots represents the fraction of cells with at least one count of the indicated gene. The color of the dots represents the average log-transformed expression of each gene across all cell types. **d**, Spatial distribution of *Fzd5* and *Wnt3* as well as hierarchical pattern of cell types. **e**, Spatial distribution pattern of cell types and marker genes within the Human Breast Cancer (Xenium) dataset. Left: Cell-type distribution through manual annotation along with four selected ROIs. Right: Distribution of *ACTA2*+ myoepithelial and endothelial cells, along with their corresponding original and imputed marker genes. **f**, tSNE plot of the cell types in Human Breast Cancer dataset. DCIS, ductal carcinoma in situ. **g**, Up-regulated genes for DCIS and invasive tumor cells in the corresponding ROIs. The size of the dots represents the -log_10_(P-values) of the differential analysis. The color of the dots represents the average log-transformed expression of each gene across four cell types. **h**, Spatial distribution of gene expression pattern of *CEACAM6, LASP1, MDM2* and *DCAF7*.

We further applied RedeFISH to the breast cancer ST dataset to investigate how tumor microenvironment changed along the transformation trajectory from ductal cells to invasive tumor cells based on the whole-transcriptome functionally-defined cells in the Xenium ST data enhanced by our algorithm. With RedeFISH results, we first identified cell types/states for the segments of the human breast cancer Xenium dataset based on the accompanied scRNA-seq data, and identified that ductal cells in different stages of transformation were found within the same Xenium slide. We selected four ROIs to capture the full spectrum of ductal cell transformation, named as DCIS #1, DCIS #2, DCIS #3 and invasive tumor cells (Fig.2e-f). Based on the whole-transcriptome expression matrix imputed by RedeFISH, we performed differential gene expression (DGE) analysis and identified specifically up-regulated genes in DCIS and invasive ROIs (Fig.2g-h). We found *CEACAM6* and *LASP1* were upregulated in DCIS cells, which have been reported to promote tumor cell migration and invasion^15, 16^, but have no significant influence on survival rate for double positive ERBB2+/ESR1+ (HER2+/ER+) breast cancer^17^ (Extended Data Fig. 12a). In contrast, *MDM2*, which plays important roles in the regulation of cellular p53 activity by a dual mechanism of degradation and transcriptional repression^18^, showed association with breast cancer prognosis (Extended Data Fig. 12a). Moreover, it has recently been reported that the over-expression of *DCAF7* is associated with poor prognosis in patients with pancreatic neuroendocrine tumors through degradation of MEN1 protein^19^. Consistently, we observed a significant association between up-regulated of *DCAF7* and survival rate in patients with breast cancer (Extended Data Fig. 12a).

In addition, we performed pseudo-time analysis based on Palantir^20^ on the RedeFISH-enhanced data, and depicted a transformation trajectory from DCIS#1, DCIS#2, DCIS#3 to invasive tumor cells (Extended Data Fig. 12b-c), consistent with the fact that DCIS is a pre-invasive form of breast cancer^21^ . As expected, DCIS #3 displayed a notably elevated median pseudotime (0.536) in comparison to DCIS #1 (0.182) or DCIS #2 (0.164), and aberrant duct morphology was indicated by the RedeFISH-enhanced data, suggestive of a transitioning state towards an invasive disease (Fig. 2e). In contrast, DCIS#1 and DCIS#2 were characterized by circular duct-like structures (Fig.2e), consistent with the pseudotime analysis results (Fig. 2g). Combining pseudotime and DGE analysis, *MDM2* and *DCAF7* exhibited significant up-regulation along the transformation trajectory (Extended Data Fig. 12d), consistent with the recent report of poor prognosis in various types of tumors ^19, 22^.

With the RedeFISH-enhanced data, we also observed several genes demonstrated pseudotemporal change of initial increase followed by decrease, such as *TENT5C*. These genes exhibited significant correlation with tumor development, thereby offering promising potential as biomarkers for investigating how tumor microenvironment changed along the transformation trajectory. We finally investigated cell-type composition in the tumor-adjacent regions and linked the cellular proportion changes with the corresponding pseudotime and the marker genes. Results suggested that transition of DCIS to invasive tumor cells was accompanied by losing surrounding myoepithelial cells and increasing vascular endothelial cells, consistent with previous observations^21^ (Extended Data Fig. 12e). In addition, we observed a decreased trend for T cell proportion, suggesting the development of immune evasion during the transition^23^ (Extended Data Fig. 12f). Moreover, more macrophages were recruited to the invasive tumor site (Extended Data Fig. 12f), probably modulating the tumor microenvironment to better accelerate the tumor progression^24^.

In summary, we develop RedeFISH, an automatic approach to identify functionally-defined cells in imaging-based ST data with whole-transcriptome expression profiles imputed thereafter. RedeFISH exhibits superior performance regarding accuracy of cell segmentation, gene imputation, and computational efficiency across various imaging-based ST platforms, without training and without the need of staining-or cell-type-label data. The cell-segmented, whole-transcriptome spatial data generated by RedeFISH enable unlocking previously infeasible investigations and generating novel biological insights.

## Acknowledgements

This work was supported by Changping Laboratory, the National Natural Science Foundation of China (32022016 X.R., 92159305 X.R., and 31991171X.R.), National Key R&D Program of China (2020YFE0202200 X.R. and 2022YFC3400904 X.R.).

## Author Contributions Statement

X.R. conceived this study, designed the algorithm, supervised the analysis, and wrote the manuscript.Y.Z. developed the software, conducted the data analysis, and wrote the manuscript.

## Methods

### Model Overview

RedeFISH employs a policy-gradient-based deep reinforcement learning approach to achieve cell alignment in imaging-based ST and scRNA-seq data by optimizing the expression similarities between ST segments and scRNA-seq cells. In detail, the pre-processing of input data is initiated to satisfy the input requirements of neural network. Then, a Segmentation Module and a Deconvolution Module are established for the purpose of carrying out region searching and alignment of single cells onto these regions. RedeFISH then proceeds to alternate between these two modules across specific epochs, yielding a collection of potential cell segments. Finally, a post-processing stage is employed to generate final polygonal regions and impute whole-transcriptome expression profiles of identified cell segments.

### Pre-processing

For data pre-processing, RedeFISH firstly encode all molecules into integers. Given an imaging-based ST data, including *n* molecules with spatial coordinates {*x*_*i*_, *y*__*i*__} and a list of genes {*g*_*i*_}, where *i*=1,…,*n*, and a *s×m* dimensions scRNA-seq expression matrix *U* with *s* cells and *m* genes. We assign a unique identification to each gene in the list *K*, which comprises *k* unique intersection genes {*k*_*i*_}, where *i*=1,…,*k*. To accomplish this, we identify genes that are present both in {*g*_*i*_} and *U*. As a result, we utilize *k* positive integer values, specifically 1, 2, …, *k*, to represent all genes in *K*. For ST data, we encode each molecule to its unique identification if *g*_*i*_ ⊆ {*k*_*i*_} and otherwise *k+*1. For scRNA-seq data, we generate a new *s×k* dimensions expression matrix *S* with *s* cells and *k* genes.

Next, in order to calculate distance scores for molecules, we implement a k-d tree structure using the Scipy package in Python to find top *d*-th nearest neighbors for each molecule based on Euclidean distance metric. Then, mean distances *d*_*i*_, *i*=1,..,*n* of the *d*-th nearest neighbors are determined and distance scores *DS*_*i*_ *i*=1,..,*n* for molecules are described as:

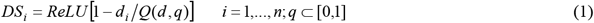

Where, *Q* is the quantile function of *d* and *ReLU* represents Rectified Linear Unit activation function. This score measures the distance between a molecule and its *d*-th nearest neighboring molecules. Hence, a molecule exhibiting a low distance score, indicating a greater seperation from other molecules, increases the probability of being classified as background noise molecules within the Segmentation Module.

### Segmentation Module

The primary objective of the Segmentation Module is to perform molecule-level classification, ultimately yielding cell regions and aggregating molecules with identical classifications to construct expression profiles. In practice, a ST dataset could contain hundreds of thousands of cells (suppose *c* cells), indicating impractical to perform

*c*-classification for each molecule when *c* is extremely large. However, given a list of candidate cells with spatial locations {*X*_*i*_, *Y*_*i*_}, *i*=1,…,*c*, RedeFISH identify top *m-*th nearest candidate cells and perform a *m*-classification for each molecule, where *m* is much smaller than *c*. The list of candidate cell locations could also be estimated by RedeFISH (see ‘Estimate Candidate Cell Locations’) or any other algorithm-based or experiment-based methods, such as clustering or staining. In addition, a *p* dimension trainable vector *f*_*j*_ represents cell features is randomly initialized for candidate cell *j*. Hence, for a pair of a molecule *i* and one of its nearest cells *j*,a feature vector *feature*_*i,j*_ is generated as:

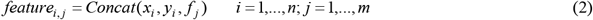

Combining molecular coordinates and cell features within feature vectors increases the probability of accurately predicting adjacent molecules belonging to the same cells.. In addition, feature vectors are further fed into a dropout layer followed by a linear layer:

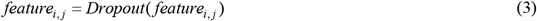

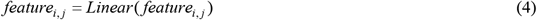

Then, softmax operation is applied to calculate assignment probabilities of molecule *i* to its *m-*th nearest cells:

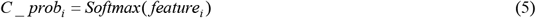

Therefore, categorical distribution *Cat*(*C_prob*_*i,1*_, *C_prob*_*i,2*_, …, *C_prob*_*i,m*_) is used to model the assignment probabilities of molecule *i* to its *m*-th nearest cells. Moreover, an additional binary classification should be performed to determine background noise molecules, thereby a trainable noise feature matrix with *c×2* dimensions is initialized in (6) followed by softmax operation in calculating probability for noise classification:

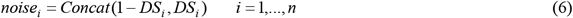

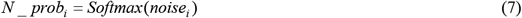

Similarly, RedeFISH utilizes categorical distribution *Cat*(*N_prob*_*i*,1_, *N_prob*_*i*,2_) to model the assignment probabilities of molecule *i* to either noise or non-noise. Based on these categorical distributions, RedeFISH samples molecule *i* to one of its nearest cells (*G*_*i*_) and either to the noise category or the non-noise category (*H*_*i*_), and then estimates the expression profile (*E*_*j,k*_) and the number of non-noise assignments (*count*_*j,e*_) for cell *j* in epoch *e* (*e*=1,…,*m_e*; *m_e* refers to max number of epochs). In addition, as a result of variations in resolution for different single-cell ST platforms, users are required to define a parameter *pc*, which represents an approximate expected number of molecules assigned to each cell. Therefore, RedeFISH calculates the probability of a given cell *j* in an epoch *e* is authentic, represented as *R*_*j,e*_, wherea higher *R* value signifies that the number of non-noise assignments is closer to the predefined *pc*. To achieve this, two normal distributions are defined as *N*(*x*|*pc, σ* ^*2*^) and *N*(*x*|0, *σ* ^*2*^), and then *R* is calculated utilizing a mixture model:

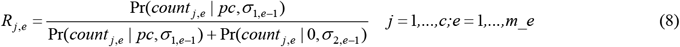

When *e* greater than 1 (not the initial epoch), the *σ* in normal distributions are calculated as:

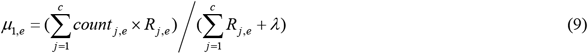

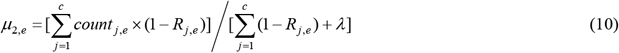

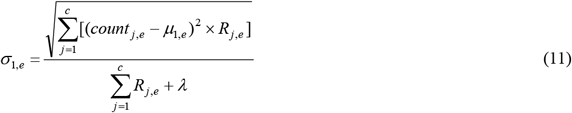

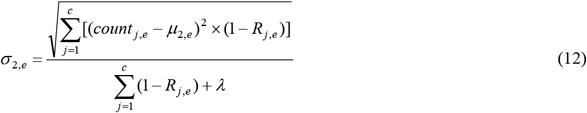

Where, *λ* is a regularization parameter. If *e* equal to 1 (the initial epoch), RedeFISH initials *σ*_*1,1*_ and *σ*_*2,1*_ as 10. Then the reward for sampling molecule *i* to cell *j* in epoch *e* is given as:

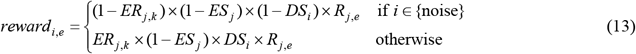

The terms *ER* and *ES* that denote expression residuals and similarities respectively will be elaborated in ‘Deconvolution Module’. Based on formula of policy gradient, we define loss function for Segmentation Module as:

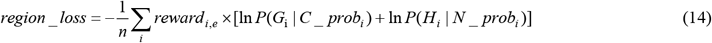

### Deconvolution Module

The primary goal of the Deconvolution Module is to improve the expression profiles of identified cells by maximizing expression similarities between cells in scRNA-seq data and the output generated by Segmentation Module. In addition, this module generates expression residuals (*ER*) and expression similarities (*ES*) that are employed in the Segmentation Module to compute rewards. Given the scRNA-seq expression matrix *S*, which comprises *t* distinct cell types, RedeFISH generates a sub-expression matrix *S*_*i*_ (*i*=1,…,*t*) with dimension of *ni×k* for each cell type, where *ni* represents number of single cells for cell-type *i*. In addition, we introduce a trainable weight variable *W*_*i*_, having dimensions of *cs×ni*, where *cs* represents the pre-defined number of cell states. Then, an expression matrix *S’* is determined:

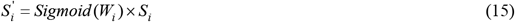

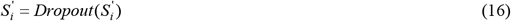

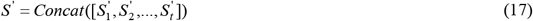

Here, the dimension of *S’* in (17) is (*cs×t*)*×k*. In practice, empolying *S’* for cell alignment not only conserves memory but also improves efficiency, as (*cs×t*) is typically significantly smaller than the total number of single cells. Also, we introduce another trainable weight variable *V* with dimensions of *c×*(*cs×t*), and this variable is applied to calculate predicted expression profiles *PE*, which is a *c×k* matrix:

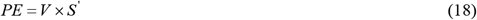

Combining *PE* with cells expression profiles *E* (in Segmentation Module), *ER* and *ES* are determined as:

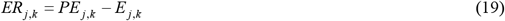

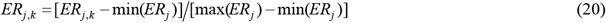

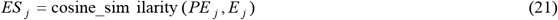

To maximize expression similarities, the loss function for the Deconvolution Module should be minimized, as represented by the following equation:

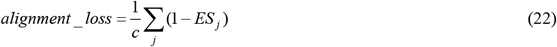

### Post-processing

Following the determination of final molecular assignments and cell expressions, post-processing is implemented to estimate polygonal regions and impute whole-transcriptome expression profiles for identified cells. In order to achieve these, RedeFISH employs alphashape in Python to create polygonal regions of cell bodies followed by fine-tuning molecular assignments based on alphashape results to generate final expression profiles. Then, a matrix *A* that represents pairwise expression cosine similarity between cells in scRNA-seq data and output of RedeFISH. For a cell *j* in the output, we identify the *r*-th most similar cells (suppose *r1,r2*,…,*rr*) from *A*_*j*_ and extract the whole-transcriptome expression profiles of these cells from *U*. Consequently, the imputed expression profile for cell *j* is computed as follows:

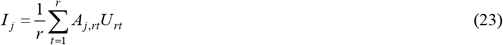

### Model Implement

At the beginning, RedeFISH preforms data pre-processing (see ‘Pre-processing’) to encode molecules in ST data and generate expression matrix of intersection genes for scRNA-seq data. To initialize the two modules, RedeFISH assigns molecules to their nearest candidate cells and generates an expression matrix for the initial assignments and the initial values of *ER* and *ES* are then determined. After that, the model undergoes iteration for a predetermined number of epochs. During each epoch, RedeFISH applies a batch training strategy to compute reward for each molecule and minimize region loss. Then, the *ER* and *ES* are subject to iterative updates for a predefined number of iterations along with minimizing alignment loss. In addition, the *ER* and *ES* obtained from previous iteration are employed to compute rewards in Segmentation Module for the next epoch. After completing all epochs, RedeFISH carries out an additional Segmentation Module to determine the final molecular assignments. In this scenario, dropout layers are omitted, and molecular assignments and noise classifications are determined by selecting the maximum probabilities within categorical distributions, rather than employing a sampling approach. In the end, RedeFISH conducts post-processing (see ‘Post-processing’) for generating polygonal regions as well as imputing whole-transcriptome expression profiles for aligned cells.

As for parameter, we set the number of epochs to 500 and initial learning rate to 0.01 and 0.001 for noise feature matrix and other variables respectively. In addition, Adam is selected as the optimizer and exponential learning rate schedule is applied with a multiplicative factor decay of 0.996 for one epoch. In pre-processing, *d* and *q* are set to 100 and 0.999 respectively for calculating the distance scores. In Segmentation Module, we set *m*=8 for cell assignments, *p*=30 for dimensions of cell features and *λ*=0.01 for regularization term. Additionally, batch size is set to the number of molecules divided by 50 with a maximum value of 2,000,000. In Deconvolution Module, we set *cs*=10 and the number of iterations in each epoch to 30. In post-processing, *r* is set to 20 for imputation of expression profiles. The manuscript consistently utilizes these default parameters in all experiments, except when specific notifications indicate otherwise.

### Estimate Candidate Cell Locations

The estimation of candidate cell locations aims at generating a list of spatial coordinates that delineates potential cell positions. To achieve this, one hypothesis is that cellular presence exhibits a positive correlation with molecular density. In addition, since cells are comprised of numerous molecules, molecular locations could serve as a reasonable approximate to represent cellular locations. Hence, we categorize molecules into either

Cell Representation Molecule (CRM) or non-CRM, and utilize the coordinates of CRMs to represent potential cell locations.

In RedeFISH, policy-gradient-based reinforcement learning is applied to determine candidate cell locations. Given an imaging-based ST data, including *n* molecules with spatial coordinates {*x*_*i*_, *y*_*i*_}, RedeFISH applies equation (1) to calculate distance score *DS*_*i*_. In addition, for molecule *i*, top *pc*-th nearest neighbors are identified (*i1,i2*,…,*ipc*) based on k-d tree and then neighbor scores *NS*_*i,h*_ are defined as:

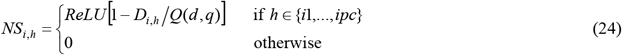

Where *D*_*i,h*_ denotes distance between molecule *i* and *h*. After that, a trainable matrix *B* with dimension of *n×*2 is initialized as *B*_*i*_ = (0,-3) to represent the feature of molecule *i* . Then, the probability for molecule *i* being classified as CRM is given as:

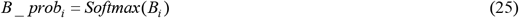

Therefore, categorical distributions *Cat*(*B_prob*_*i*,1_, *B_prob*_*i*,2_) are employed for sampling molecule *i* to either CRM or non-CRM. In addition, a coverage vector *CO* with a length of *n* is applied for calculating the coverage score *CS*.

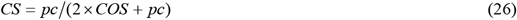

Where, *COS*_*i*_ refers to sum *CO* value of *pc*-th nearest neighbors (suppose *CO*_*i*1_,…,*CO*_*ipc*_) of molecule *i*.

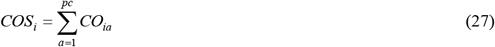

The reward and loss function are defined as:

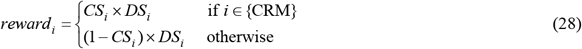

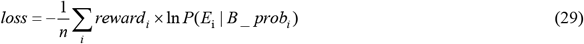

RedeFISH utilizes an Adam optimization algorithm with an initial learning rate of *n*\*10^−10^ and an exponential learning rate schedule to minimize the loss function. As default, we train the above model for 1000 epochs and 100 iterations per epoch.

At the beginning of each epoch, values in *CO* are initialized to zero. During each iteration, RedeFISH selects *n*/100 non-redundant molecules and classifies them as CRM or non-CRM followed by minimizing loss function. Additionally, *CO* is updated at the end of each iterative, with the underlying objective for computing reward and loss in the next iteration. Suppose set *G* refers to all CRMs in the iteration, then *CO* is updated by:

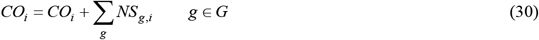

After completing the 1000 epochs, an additional epoch is carried out for determining final CRMs through selection of maximum probabilities within categorical distributions. Therefore, a list of CRMs’ coordinates is generated to represent candidate cell locations.

## Datasets for Benchmarking

*Human Non-small Cell Lung Cancer*. ST dataset of Human non-small cell lung cancer (NSCLC) from CosMx platform is available at https://nanostring.com/products/cosmx-spatial-molecular-imager/nsclc-ffpe-dataset/^4^. We chose Lung5_Rep1 in this study, including 37,226,610 transcripts for 980 genes. Meanwhile, an h5ad file containing annotation information pertaining to scRNA-seq dataset is accessible at

https://cellxgene.cziscience.com/collections/edb893ee-4066-4128-9aec-5eb2b03f8287^25^.

*Human Breast Cancer*. Replicate 1 of human breast cancer ST dataset is available at https://www.10xgenomics.com/products/xenium-in-situ/preview-dataset-human-breastbased on Xenium platform^5^, including 43,664,530 transcripts for 541 genes. In addition, scFFPE-seq dataset from the same sources was applied for RedeFISH algorithm. We used the clustering result to perform RedeFISH, since annotation is not available.

*Mouse Ileum*. The MERFISH based mouse ileum dataset is available at https://doi.org/10.5061/dryad.jm63xsjb2 ^10^ comprised of 819,665 transcripts for 241 genes. We additionally downloaded scRNA-seq dataset at https://singlecell.broadinstitute.org/single_cell with an accession of SCP1038^26^. Moreover, cell annotation information was extracted from corresponding metadata file.

*Mouse Brain MERFISH*. We downloaded sample S1R1 MERFISH dataset of mouse brain from Vizgen website (https://vizgen.com/data-release-program/) ^27^, including 54,712,414 for 649 genes. Additionally, snRNA-seq dataset, accompanied by relevant annotations, was downloaded from

https://www.ebi.ac.uk/arrayexpress/experiments/E-MTAB-11115/ ^28^.

*Mouse Liver*. Sample L1R2 MERFISH dataset of mouse liver is available at https://vizgen.com/data-release-program/ ^29^ comprised of 225,890,863 transcripts for 385 genes. We obtained droplet scRNA-seq dataset with accompanying annotations from https://figshare.com/articles/dataset/Processed_files_to_use_with_scanpy_/8273102/2 ^30^.

*Mouse Brain Stereo-seq*. The Stereo-seq based mouse brain ST dataset was downloaded from MOSTA website (https://db.cngb.org/stomics/mosta/download/) ^11^. In addition, We procured scRNA-seq dataset and corresponding metadata from Allen Brain Atlas (https://portal.brain-map.org/atlases-and-data/rnaseq/mouse-whole-cortex-and-hippocampus-10x) ^31^.

### Implement RedeFISH on Datasets

We implemented RedeFISH on the aforementioned datasets using the default setting, with the exception of any special considerations for each dataset. For Human-NSCLC dataset, we used the annotation from “cell_type_major” in the h5ad file that includes 24 major cell types. For Human Breast Cancer dataset, we labeled each cluster in scFFPE-seq data with a corresponding cell-type designation (Extended Data Table. 1). For Mouse Ileum dataset, we relabeled 3 subclasses of transit-amplifying (TA) cells (“TA_3”, “TA_4” and “TA_5”) to enterocyte cells, because they highly expressed *Slc51a* and *Slc5a1*, the markers of enterocyte cells. In addition, we set *q* to 0.9999 in data pre-processing. For mouse liver dataset, we excluded Hepatocyte (Pericentral, Midlobular, Periportal) and Hepatocyte (Pericentral and Periportal) cells due to ambiguity regarding their precise spatial distribution. Finally, *pc* was set to 200, 160, 100, 400 and 600 for Human-NSCLC, Human Breast Cancer, Mouse Ileum, Mouse Brain (MERFISH) and Mouse Liver datasets respectively in order to keep the number of identified cells roughly equal to the number from originally published segmentation.

As for sequencing-based ST platforms, such as Stereo-seq, an additional pre-processing procedure was conducted. We initially derived a list of molecules with spatial coordinates by transforming the spot-by-gene expression matrix into an imaging-based format, wherein each sequencing-based unique molecular identifier (UMI) was treated as a discrete molecule. Then, top *k* intersection genes, exhibiting the highest levels of expression in ST data, were identified and subsequently encoded as positive integers from 1 to *k*, whereas other genes were encoded as *k*+1. In addition, the spot-over-count (*soc*) metric was computed as the ratio of the total number of spots to the total number of molecules. We adapted the aforementioned method and implemented RedeFISH on Mouse Brain (Stereo-seq) dataset. The *k* was set to 2000 for intersection genes and *q* was set to 0.9999 in calculating distance score. Moreover, we set *pc* equal to 2000\**soc* for estimating candidate cell locations.

### Implement Baysor on Datasets

We implemented Baysor v0.5.0 ^10^ on the benchmarking datasets. Firstly, we used “--save-polygons GeoJSON” to generate polygonal regions of segmented cells. Then the scale parameter was set to 50, 7, 30 and 9 for Human-NSCLC (CosMx), Human Breast Cancer (Xenium), Mouse Ileum (MERFISH), Mouse Brain (MERFISH) datasets respectively to maintain an approximately equivalent cellular radius between segmented cells from Baysor or original publication. In addition, the implementation of Baysor is infeasible for ultra-large datasets, such as Mouse Liver and Mouse Brain (Stereo-seq) datasets, due to memory limitation (exceed 5TB).

### Expression Similarity

As one goal of RedeFISH is to optimize expression profiles of aligned cells, we measured expression similarity between aligned cells and single cells in scRNA-seq data. To achieve this, cosine similarity and root mean square error (RMSE) were calculated as evaluation metrics for assessing the similarities. We firstly identified intersection genes between ST and scRNA-seq data and generated expression matrix *A* and *B* based on the shared genes. The total UMI counts of each cell in *A* and *B* was additionally normalized to the total number of intersection genes. Thus, given a aligned cell *i* and single cell *j*, the corresponding cosine similarity (*SIM_COS*_*i,j*_) and RMSE (*SIM_RMSE*_*i,j*_) were computed as:

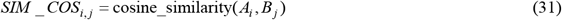

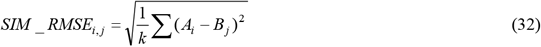

Where, *k* refers to number of intersection genes. For each aligned cell *i*, we generated two lists of similarities between *i* and all single cells, therefore {*SIM_COS*_*i*,1_,…,*SIM_COS*_*i,n*_} and {*SIM_RMSE*_*i*,1_,…,*SIM_RMSE*_*i,n*_}. Finally, overall similarities *SIM_C*_*i*_ and *SIM_R*_*i*_ were calculated by averaging top 20 highest values in the first list and top 20 lowest values in the second list respectively. We compared *SIM_C* and *SIM_R* for all identified cells for the three approaches based on boxplot across six datasets. A higher *SIM_C* and lower *SIM_R* signfy an improved level of expression similarity between aligned cells and single cells. The corresponding scRNA-seq data for similarities calculation in this section were given in “Datasets for Benchmarking” section.

To pair-wisely compared expression similarities of identified cells in alternative approaches. We built a uniform four-connected grid over 2D space, maintaining a Euclidean distance of *d*/3 between each pair of adjacent grid nodes. Then, we assigned one identified cell to each grid node based on spatial proximity and employed the expression profile of identified cells to represent the expression of the corresponding grid node. In addition, If there were no identified cells located within a distance of 3\**d* from a given grid node, the expression value of zero was attributed to that particular grid node. We calculated expression similarities between single cells and grid nodes that have non-zero expression in a minimum of two approaches based on formula (31) and (32). The values of *d* were set to approximately match the cell radius, specifically 50, 7, 30, 9 and 9 for Human-NSCLC (CoxMx), Human Breast Cancer (Xenium), Mouse Ileum (MERFISH), Mouse Brain (MERFISH) and Mouse Liver (MERFISH) datasets respectively

### Label Transferring

To facilitate comparison of cell-type distributions, label transferring was applied to automatically assign cell-type from scRNA-seq data to identified cells. Given an identified cell *i*, we firstly selected top 20 cells from the scRNA-seq data (suppose *O*_*i*_) that exhibit the highest cosine similarity of expression profile with the cell.. In addition, supposing CT(*o*) represents cell-type of cell *o*, we calculated probability of cell *i* belong to cell-type *k* as:

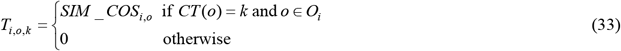

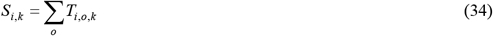

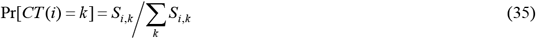

Hence, we predicted cell-type for all identified cells based on the probabilities.

### Runtime Efficiency

The computational efficiencies were assessed by comparing total execution times for RedeFISH and Baysor. We tested the run time of RedeFISH on a single NVIDIA A100 card. However, due to absence of GPU acceleration support, we executed Baysor on a computer with Intel(R) Xeon(R) Platinum 8253 CPU.

### Analyze Mouse Ileum Dataset

We applied Leiden for cell clustering based on the expression profiles of identified cells, followed by manual cell annotation using marker genes. Then, tSNE in Scanpy packages was applied for cell-type illustration. In addition, CellPhoneDB ^32^ was employed for calculating significant of interaction between *Wnt* and *Fzd-Lrp* complexes.

### Analyze Human Breast Cancer Dataset

Initially, Leiden combined with manual annotation was employed for the purpose of cell-type annotation. Based on the spatial distribution of cell types, four regions of interests (ROIs) were selected for downstream analysis, namely DCIS #1, DCIS #2, DCIS #3 and invasive tumor. To investigate up-regulated genes in ROIs, we randomly selected 300 cells from one ROI and 100 cells from each of the remaining three ROIs, which was designated as the control group. Then, edgeR ^33^ was applied to identify up-regulated genes and compute the level of statistical significance. In addition, we used GEPIA ^17^ to analyze survival rate for several genes. In this analysis, we chose BRCA HER2+ non-luminal dataset, because the tissue block was annotated as HER2+/ER+/PR− ^5^, and set 60% and 40% for parameter Cutoff-High and Cutoff-Low respectively. For pseudotime analysis, we randomly chose one start cell and three terminal cells from DCIS #1 and invasive tumor ROIs respectively, and applied Palantir ^20^ with default setting for estimating pseudotime for DCIS and invasive tumor cells. We further employed a smooth spline method to perform curve fitting between pseudotime and gene expressions or cell-type proportions.

To investigate cell-type proportions in tumor microenvironment, we firstly generated a list of spatial coordinates containing all cells with indicated cell type in each ROI. For example, a list of coordinates of DCIS #1 cells in DCIS #1 ROI. We then identified all cells whose Euclidean distance from any one of the coordinates in the list falls within a specified radius (10μm, 20μm, 50μm and 100μm), which enabled us for the proportion calculation. Hence, we calculated the cell-type proportions of these cells for each DCIS or invasive tumor cells.

## Data availability

*Human-NSCLC*. The ST dataset is available at

https://nanostring.com/products/cosmx-spatial-molecular-imager/nsclc-ffpe-dataset/. The scRNA-seq dataset with annotation is available at https://cellxgene.cziscience.com/collections/edb893ee-4066-4128-9aec-5eb2b03f8287.

*Human Breast Cancer*. Both ST and scFFPE-seq dataset of human breast cancer are available at https://www.10xgenomics.com/products/xenium-in-situ/preview-dataset-human-breast.

*Mouse Ileum*. The mouse ileum ST dataset is available at https://doi.org/10.5061/dryad.jm63xsjb2. The scRNA-data is available at https://singlecell.broadinstitute.org/single_cell with an accession of SCP1038.

*Mouse Brain (MERFISH)*. The mouse brain (MERFISH) dataset is available at https://vizgen.com/data-release-program/. The snRNA-seq dataset with annotation is available at https://www.ebi.ac.uk/arrayexpress/experiments/E-MTAB-11115/.

*Mouse Liver*. The mouse liver (MERFISH) dataset is available at https://vizgen.com/data-release-program/. The droplet scRNA-seq dataset with annotation is available at https://figshare.com/articles/dataset/Processed_files_to_use_with_scanpy_/8273102/2.

*Mouse Brain (Stereo-seq)*. The mouse brain (Stereo-seq) dataset is available on MOSTA website (https://db.cngb.org/stomics/mosta/download/). The scRNA-seq dataset and corresponding metadata is available on Allen Brain Atlas (https://portal.brain-map.org/atlases-and-data/rnaseq/mouse-whole-cortex-and-hippocampus-10x).

## Code availability

The package is available on github with detailed documentation (https://github.com/Roshan1992/RedeFISH).

---

**Extended Data Fig. 1.**
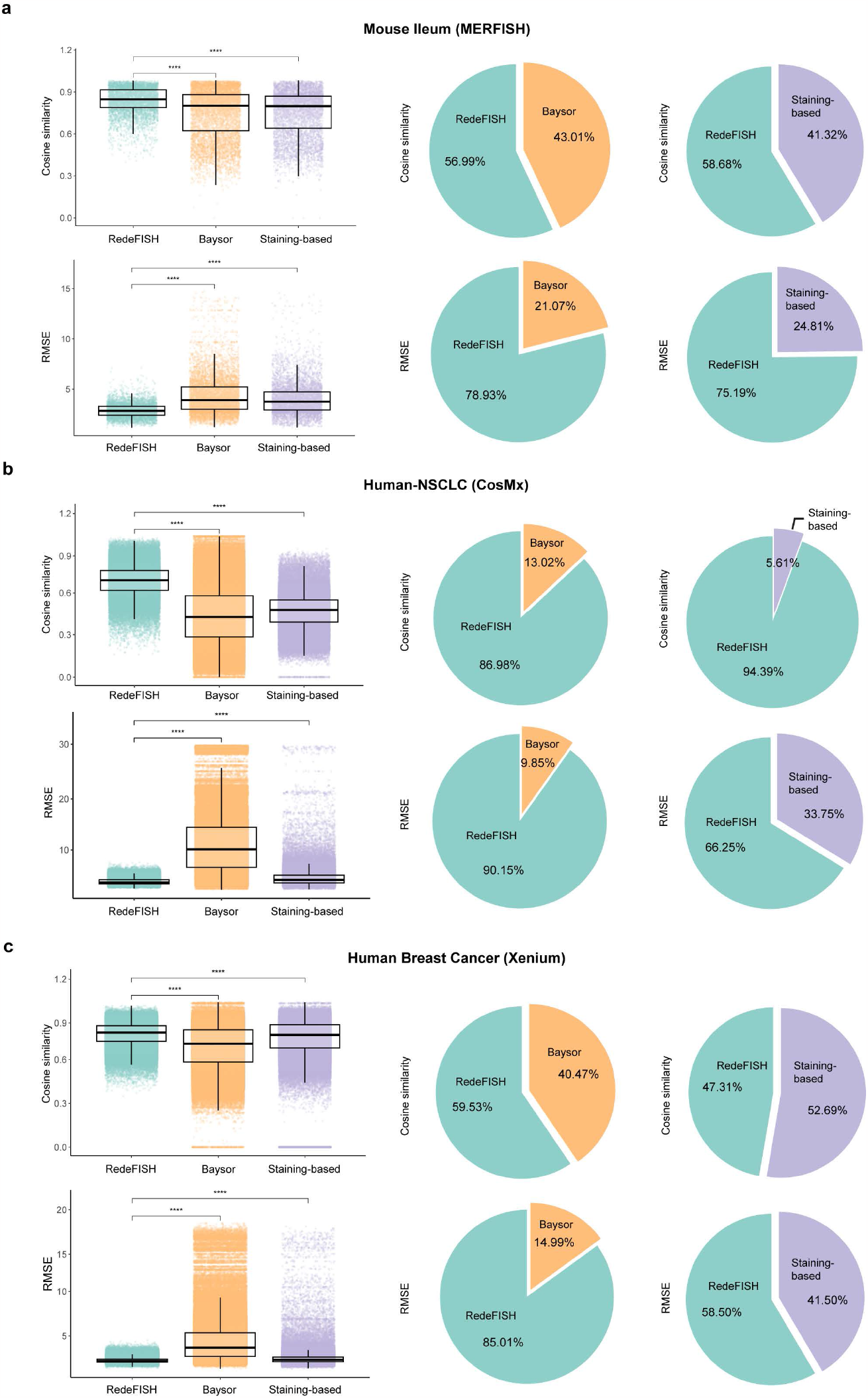
Expression similarity between identified cells and single cells on Mouse Ileum (MERFISH), Human-NSCLC (CosMx) and Human Breast Cancer (Xenium) datasets. **a-c**, Cosine similarity and RMSE of expression profiles between single cells in scRNA-seq data and outputs from RedeFISH, Baysor, and Staining-based method of the three datasets. Left: Boxplot of cosine similarity or RMSE for identidied cells. Right: Pie charts depict the proportion of grid nodes that exhibit superior expression similarity for RedeFISH or alternative approaches.

**Extended Data Fig. 2.**
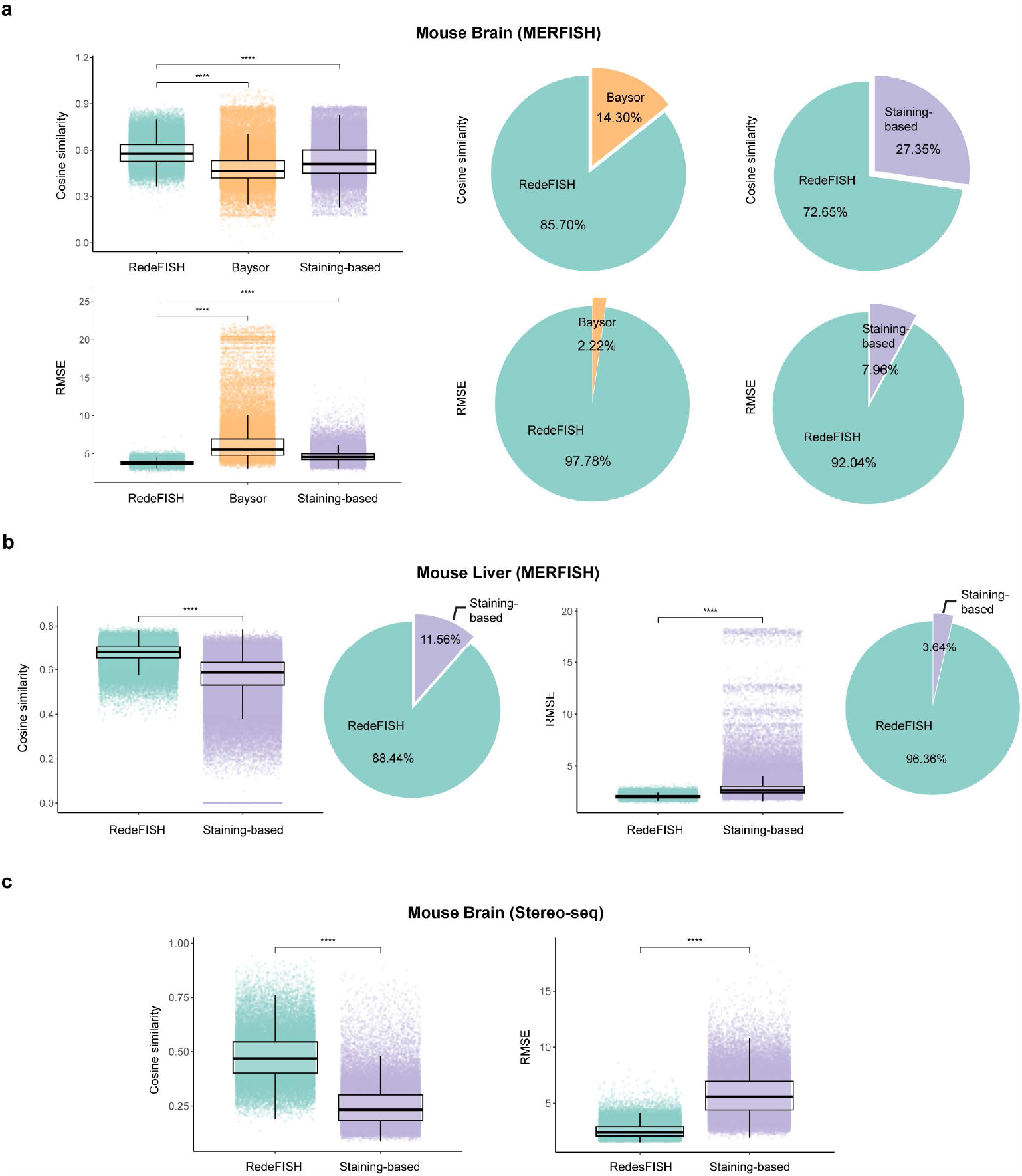
Expression similarity between identified cells and single cells on Mouse Brain (MERFISH), Mouse Liver (MERFISH) and Mouse Brain (Stereo-seq) datasets. **a-c**, Cosine similarity and RMSE of expression profiles between single cells in scRNA-seq data and outputs from RedeFISH, Baysor, and Staining-based method of the three datasets. Boxplot: Boxplot of cosine similarity or RMSE for identidied cells. Pie charts: Pie charts depict the proportion of grid nodes that exhibit superior expression similarity for RedeFISH or alternative approaches.

**Extended Data Fig. 3.**
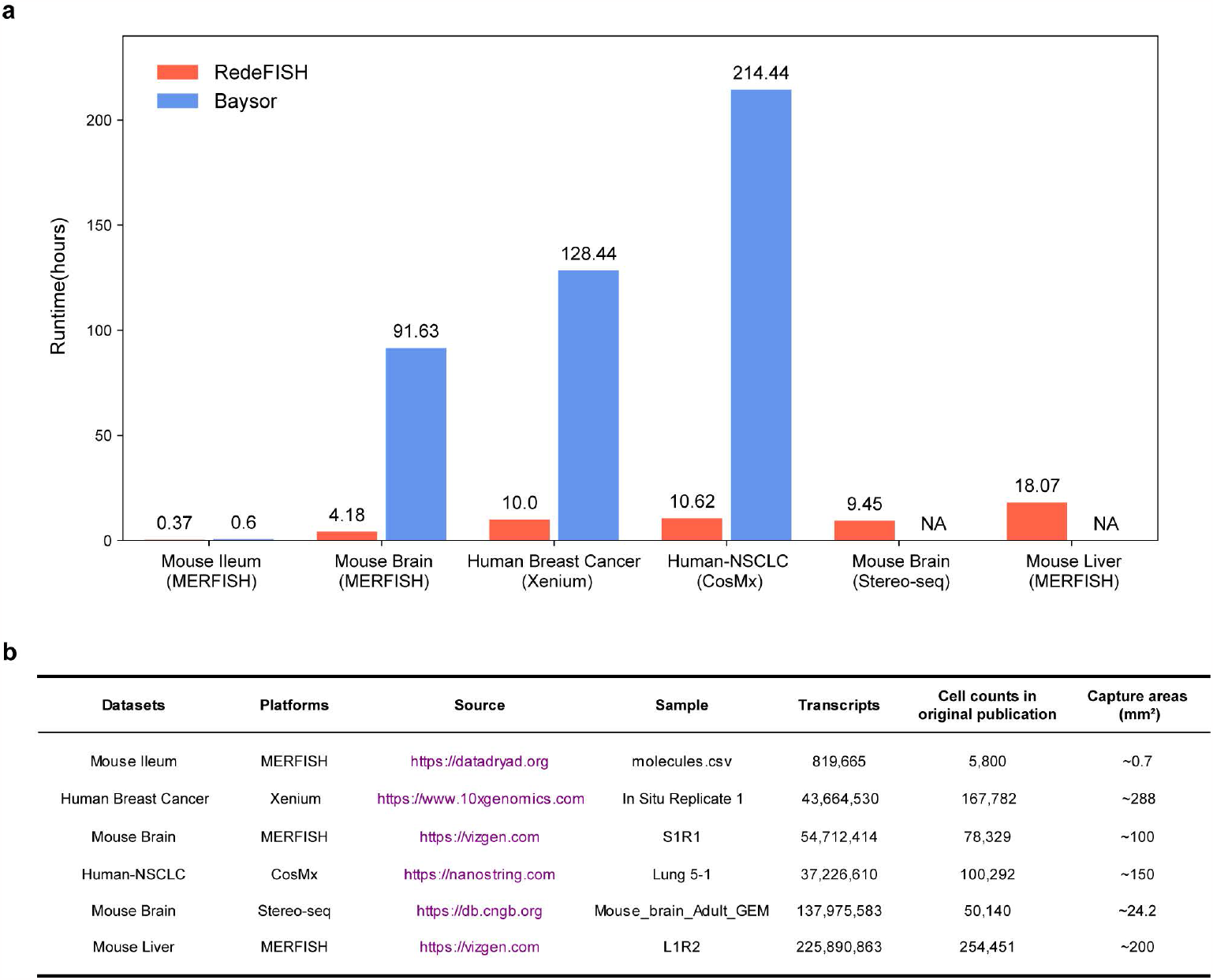
Runtime efficiency of RedeFISH and Baysor. **a**, Comparison of run time (hours) between RedeFISH and Baysor on the six datasets. **b**, Table of detail information includes platforms, source, sample, number of transcripts and cell counts in original publication of the six datasets.

**Extended Data Fig. 4.**
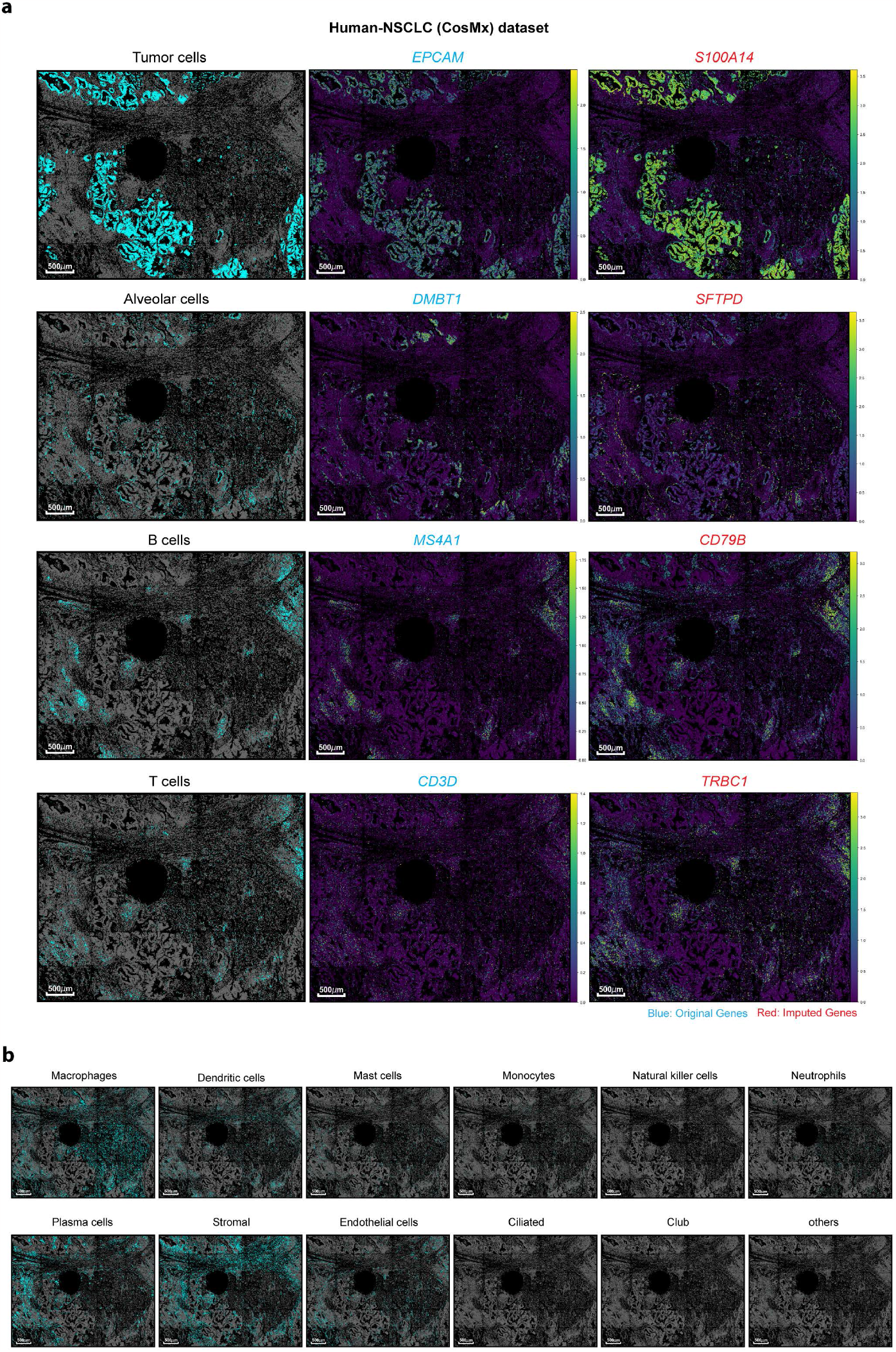
Spatial distributions of all cell types by RedeFISH on Human-NSCLC (CosMx) dataset through label transferring. **a**, Spatial distribution of tumor, alveolar, B and T cells along with their corresponding original and imputed marker genes. **b**, Spatial distribution 12 cell types. Gray dots represent the location of all cells and cyan dots represent the location of the indicated cell type.

**Extended Data Fig. 5.**
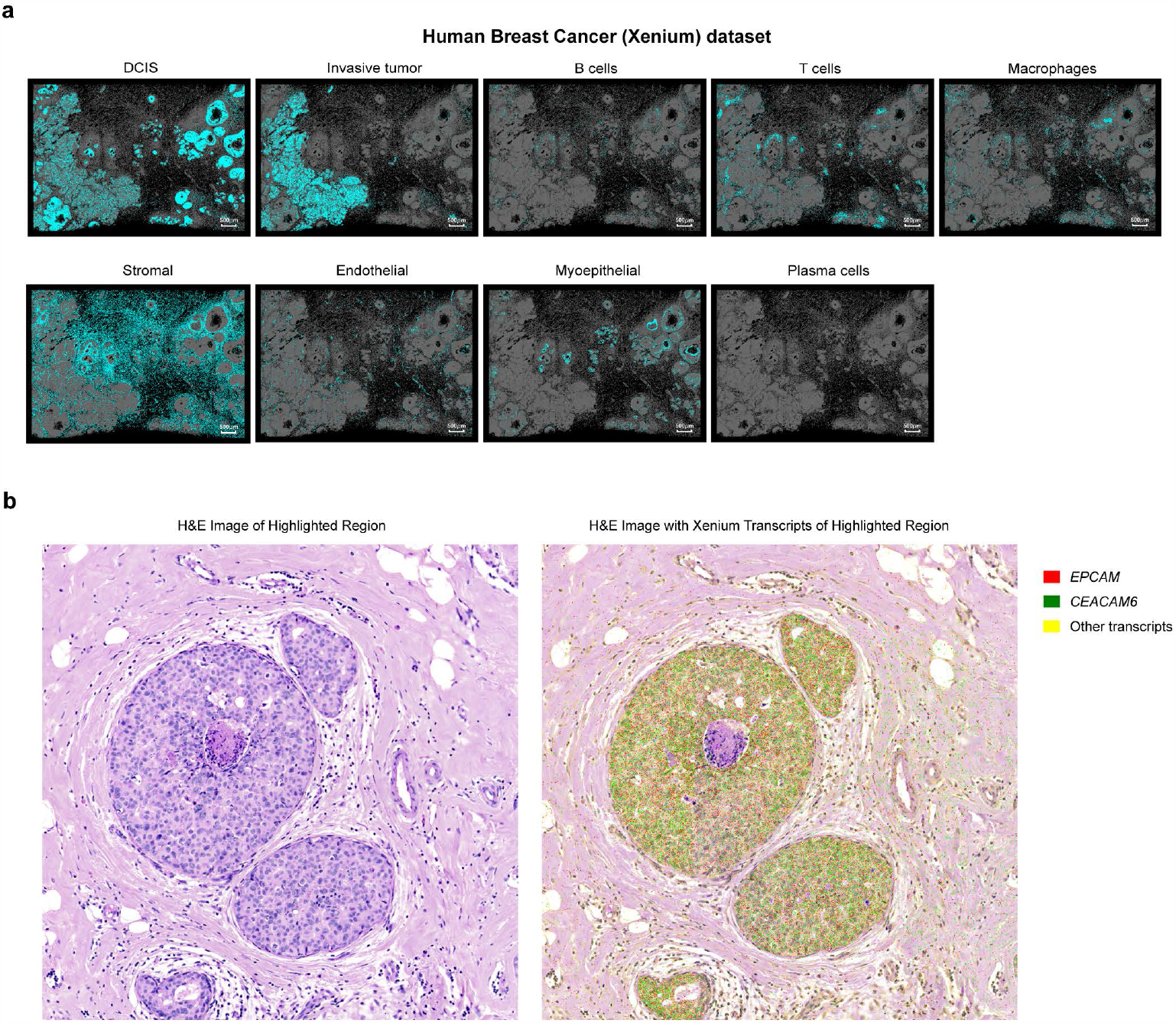
Spatial distribution of cell types, H&E image and Xenium trainscripts on Human Breast Cancer (Xenium) dataset. **a**, Spatial distributions of 9 cell types by RedeFISH through label transferring. Gray dots represent the location of all cells and cyan dots represent the location of the indicated cell type. DCIS, ductal carcinoma in situ. **b**, H&E image and Xenium transcripts of the highlighted region.

**Extended Data Fig. 6.**
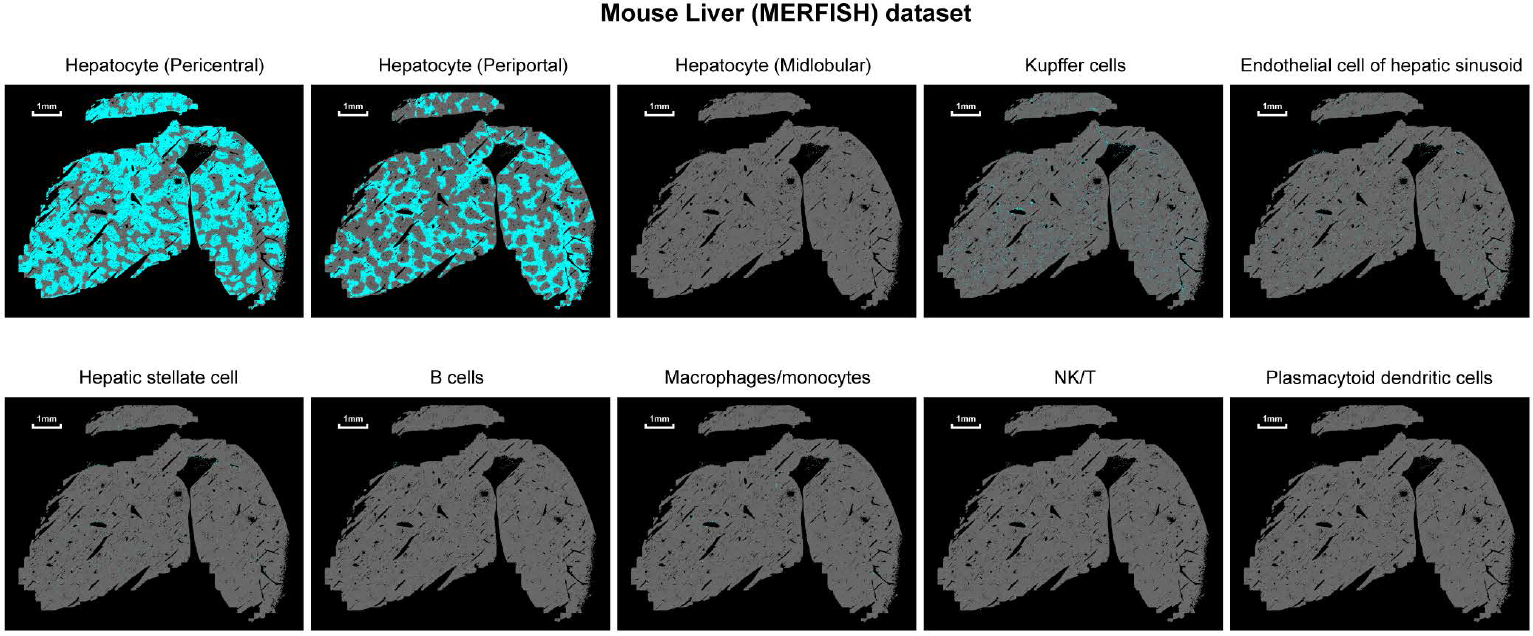
Spatial distributions of all cell types by RedeFISH on Mouse Liver (MERFISH) dataset through label transferring. Gray dots represent the location of all cells and cyan dots represent the location of the indicated cell type. NK, natural killer.

**Extended Data Fig. 7.**
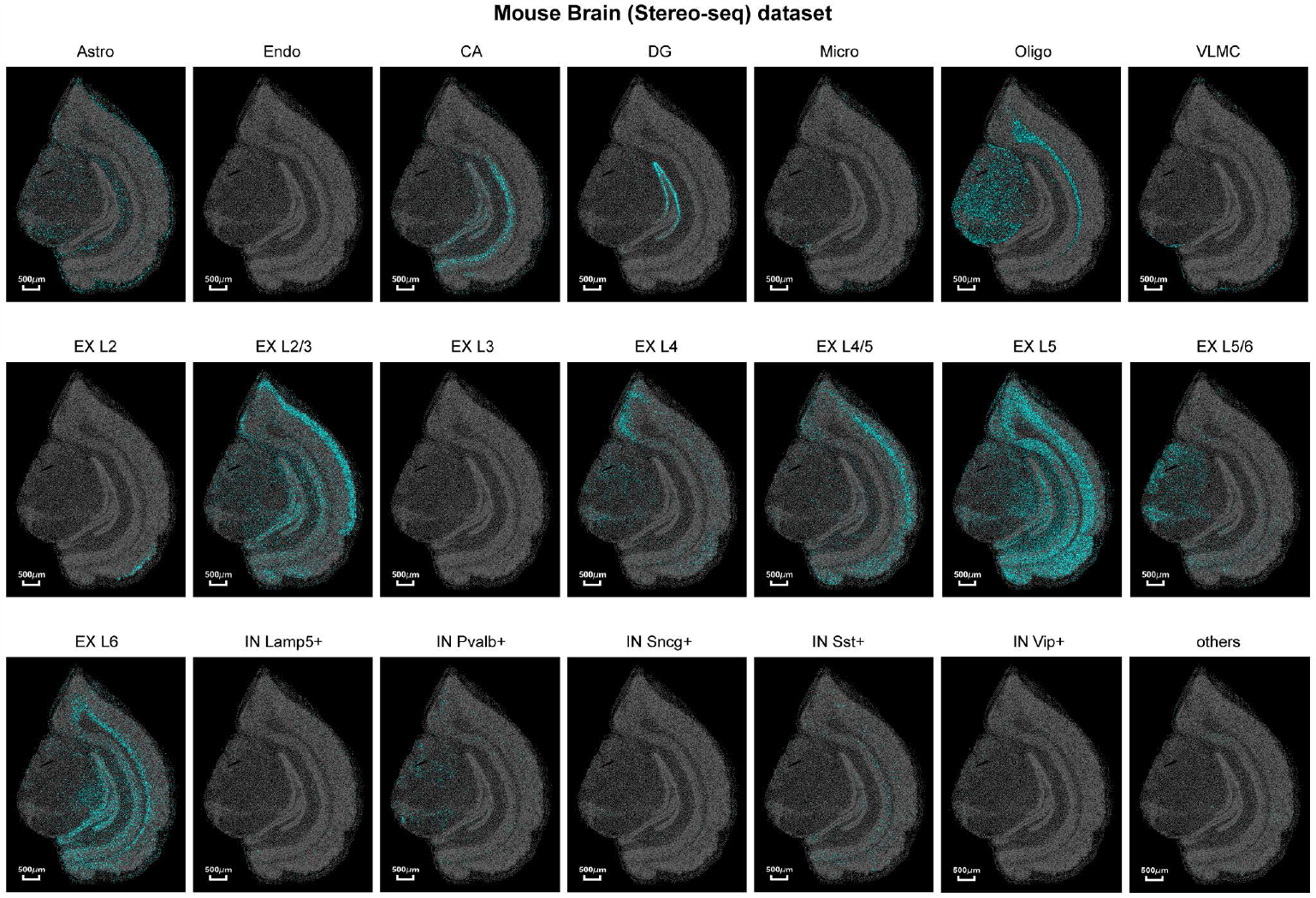
Spatial distributions of all cell types by RedeFISH on Mouse Brain (Stereo-seq) dataset through label transferring. Gray dots represent the location of all cells and cyan dots represent the location of the indicated cell type. Astro, astrocyte; CA, cornu ammonis area; DG, dentate gyrus; Endo, endothelial cell; EX, excitatory glutamatergic neuron; IN, GABAergic interneuron; Micro, microglia; Oligo, oligodendrocyte; VLMC, vascular and leptomeningeal cells.

**Extended Data Fig. 8.**
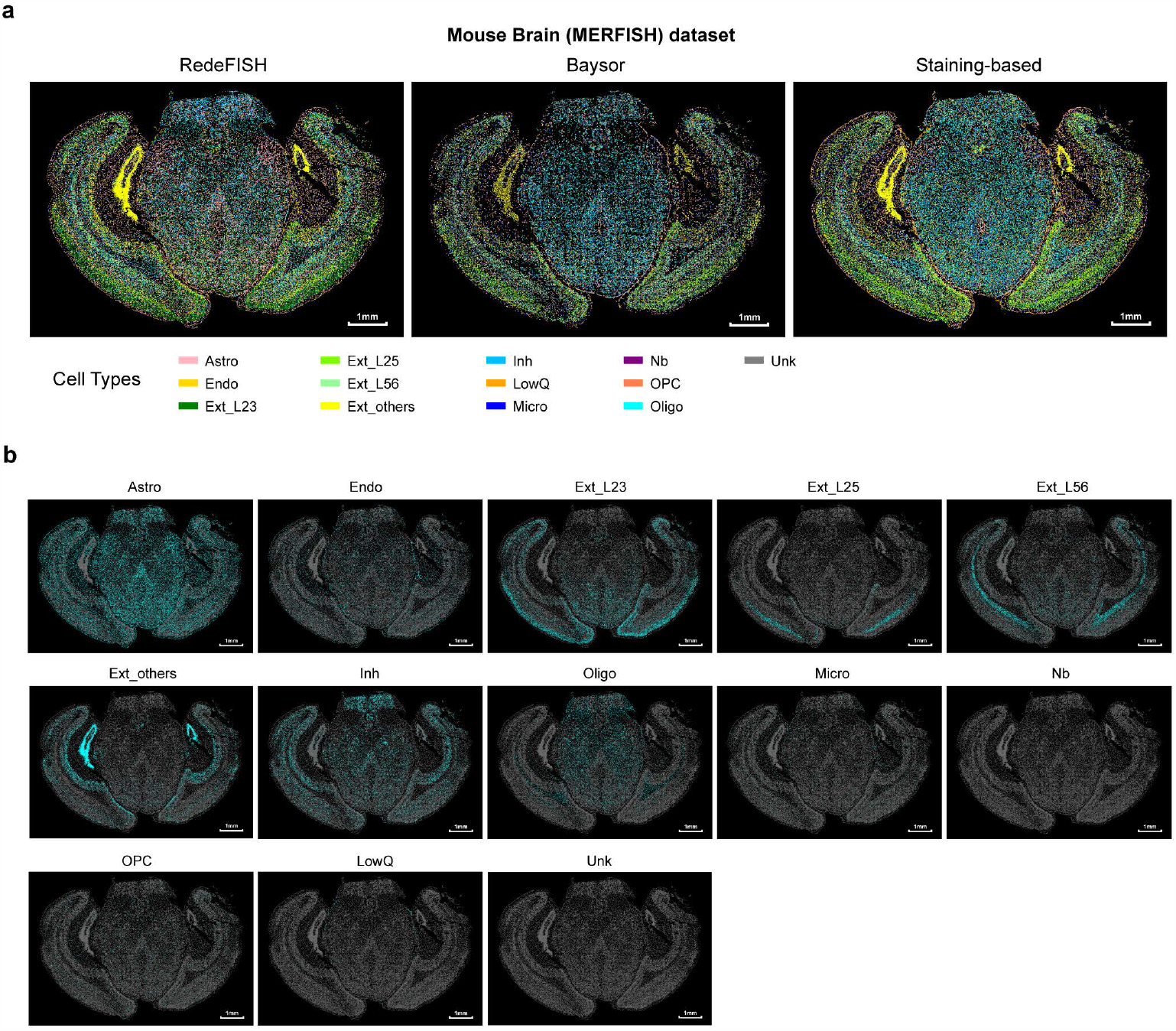
Spatial distributions of all cell types on Mouse Brain (MERFISH) dataset through label transferring. **a**, Cell-type distribution of all identified cells from RedeFISH, Baysor and Staining-based method. **b**, Distribution of indicated cell types based on RedeFISH results. Gray dots represent the location of all cells and cyan dots represent the location of the indicated cell type. Astro, astrocyte; Endo, endothelial cell; Ext, excitatory neurons; Inh, thalamic habenular neuron; LowQ, low quality cells; Micro, microglia; Oligo, oligodendrocyte; OPC, oligodendrocyte precursor cell; Unk, Unknow.

**Extended Data Fig. 9.**
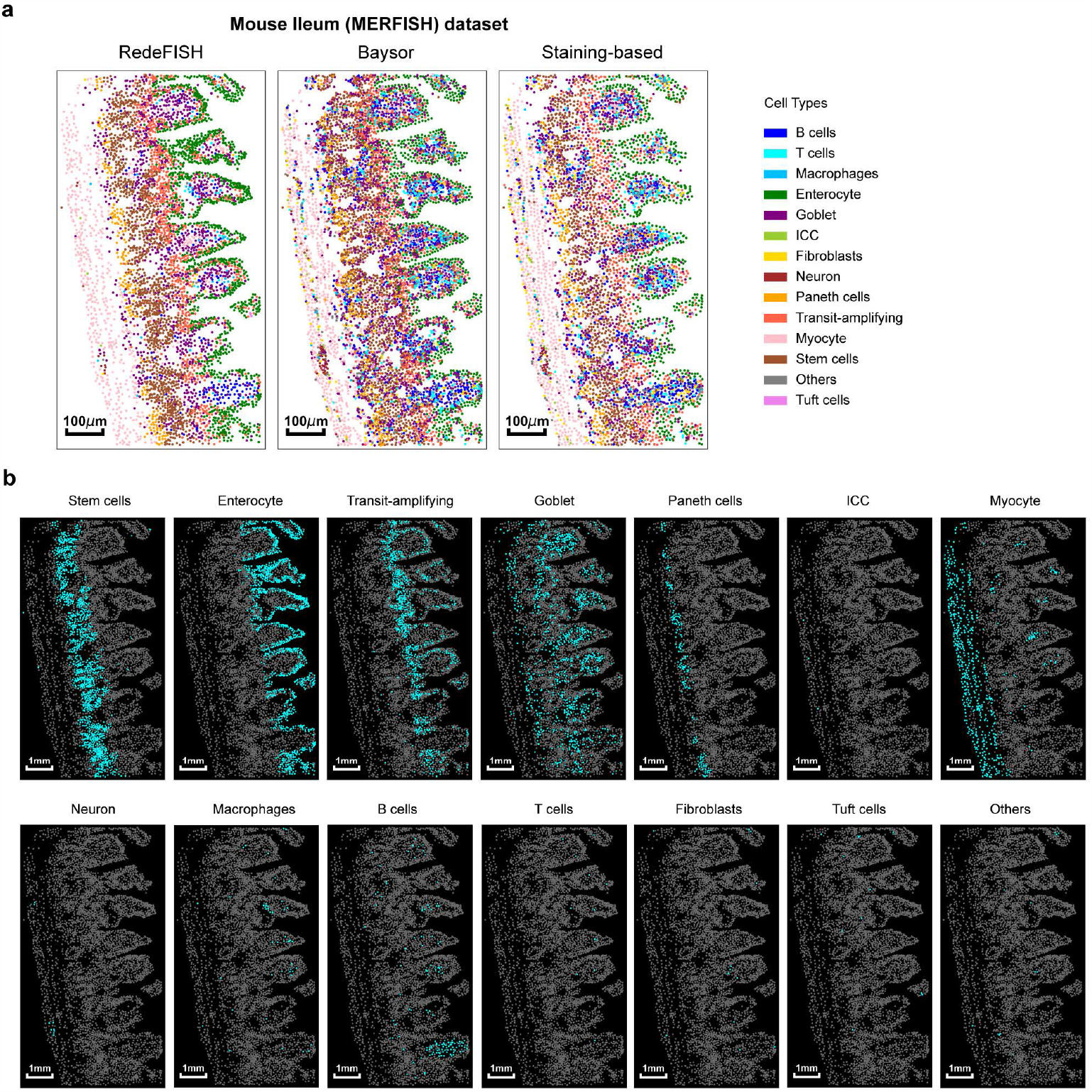
Spatial distributions of all cell types on Mouse Ileum (MERFISH) dataset through label transferring. **a**, Cell-type distribution of all identified cells from RedeFISH, Baysor and Staining-based method. **b**, Distribution of indicated cell types based on RedeFISH results. Gray dots represent the location of all cells and cyan dots represent the location of the indicated cell type. ICC, Interstitial cells of Cajal.

**Extended Data Fig. 10.**
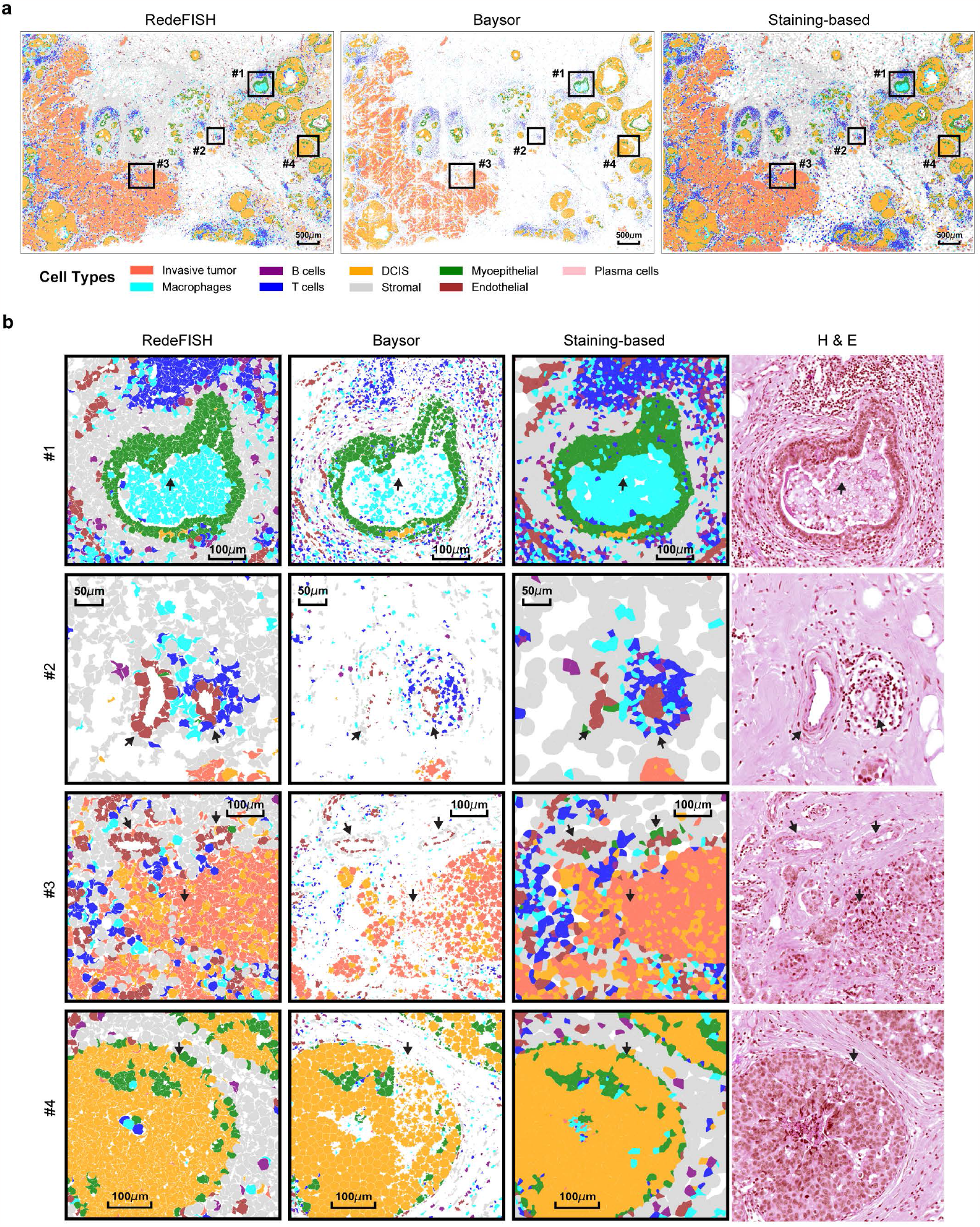
Regions of identified cells on Human Breast Cancer (Xenium) dataset. **a**, Distribution of cell bodies colored by cell types for the results obtained from RedeFISH, Baysor, and Staining-based method along with four selected ROIs. **b**, Distribution of cell bodies in each ROI for RedeFISH, Baysor, and Staining-based method combined with H&E image of corresponding regions.

**Extended Data Fig. 11.**
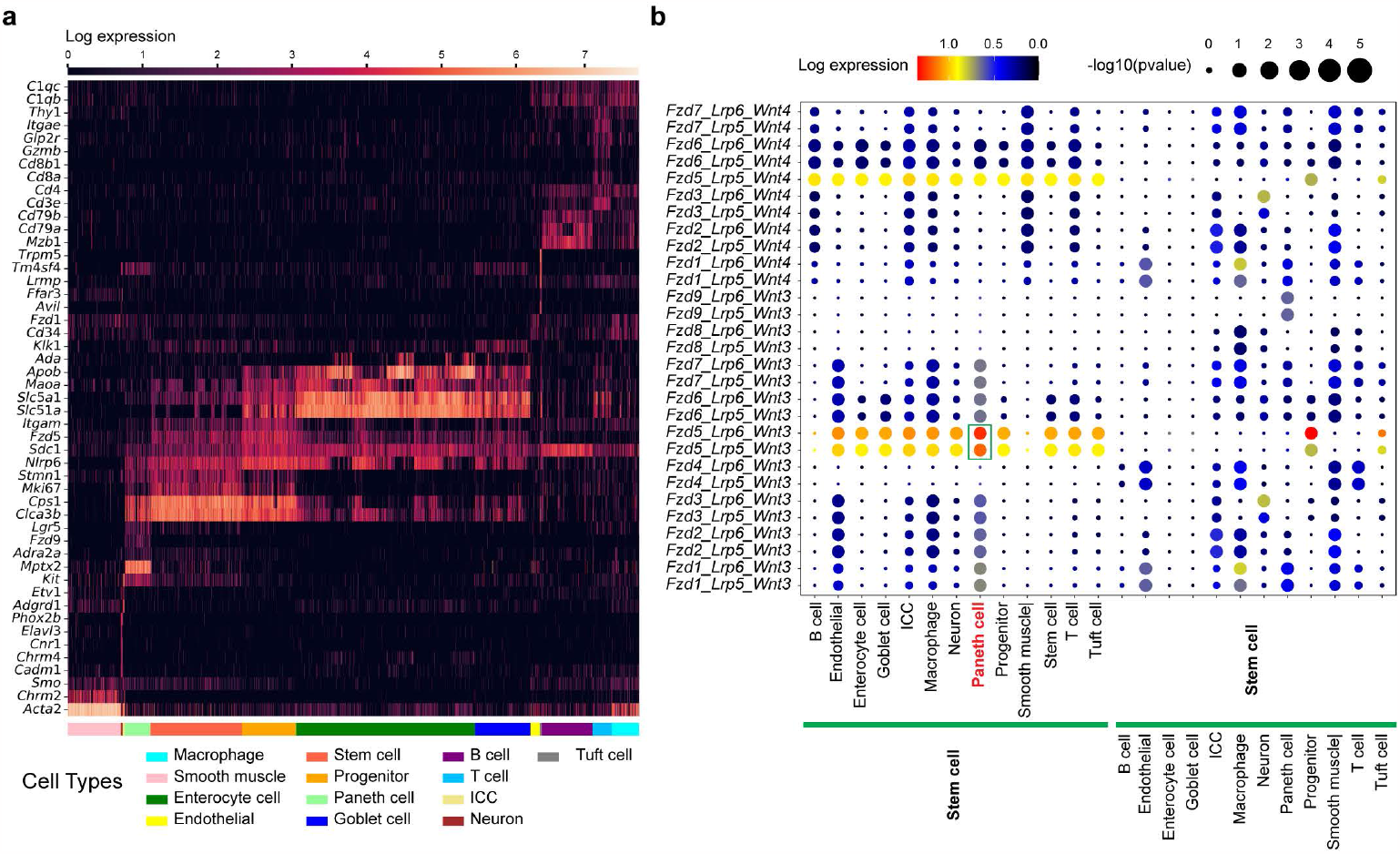
Analysis of RedeFISH results from Mouse Ileum (MERFISH) dataset. **a**, Cell expression of marker genes for cell types. **b**, Expression of *Wnt* and *Fzd-Lrp* complexes between stem cell and various cell types. The size of the dots represents the -log_10_(P-values) of the interaction significance. The color of the dots represents the average log-transformed expression of ligand and receptor.

**Extended Data Fig. 12.**
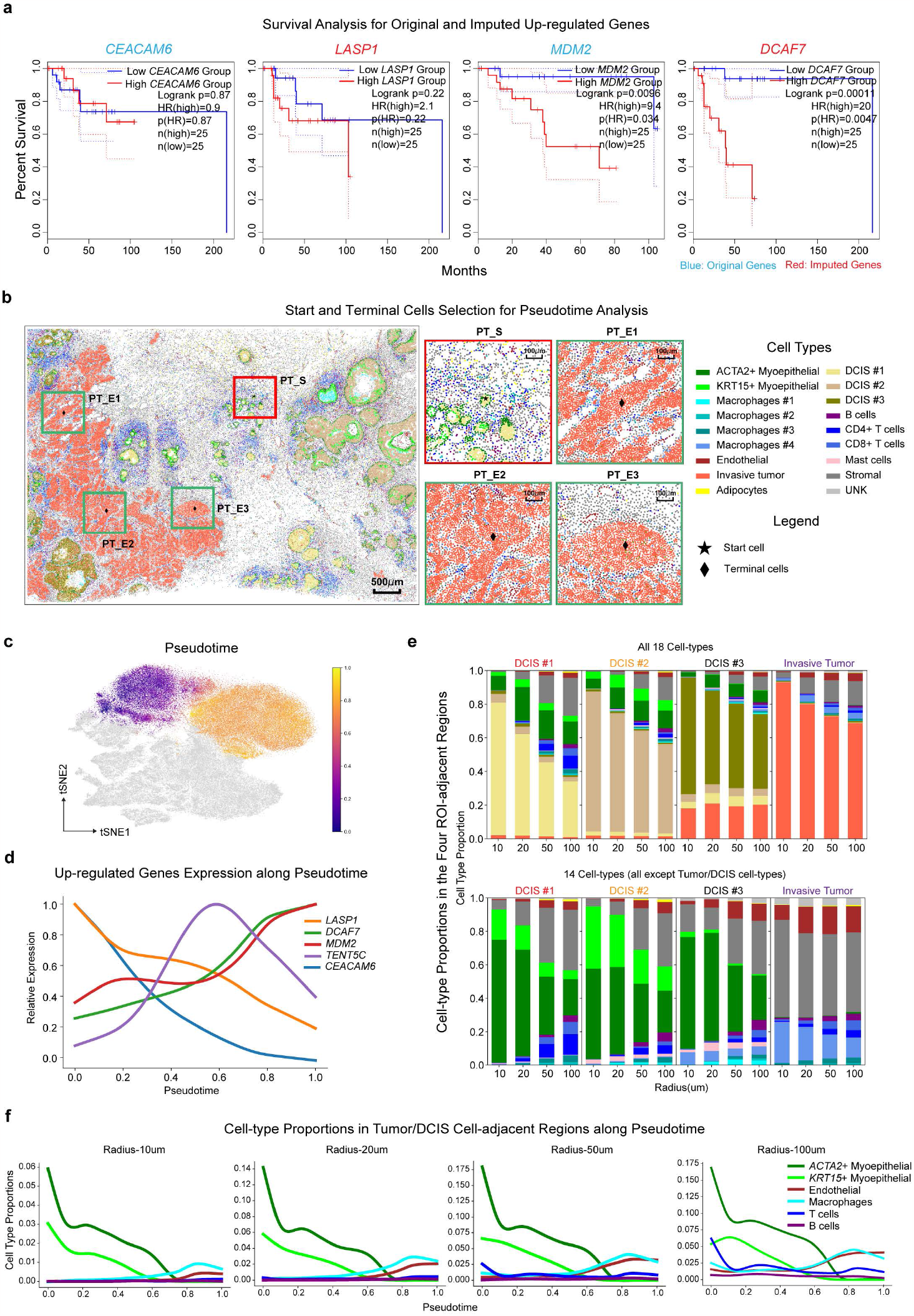
Analysis of RedeFISH results from Human Breast Cancer (Xenium) dataset. **a**, Survival curves for *CEACAM6, LASP1, MDM2* and *DCAF7*. **b**, The start cell and terminal cells from DCIS #1 and invasive tumor ROIs for analyzing pseudotime. **c**, Pseudotime of DCIS and invasive tumor cells on the tSNE coordinates. **d**, Line chart showing the normalized expression of five selected genes along pseudotime. **e**, Percentage of cell types found in neighboring areas of DCIS or invasive tumor cells. Top: Percentages for 18 cell types. Button: Percentages for 14 cell types excluding DCIS #1, DCIS #2, DCIS #3 and invasive tumor. **f**. Line chart depicting the proportions of six major cell types in regions located within 10μm, 20μm, 50μm, and 100μm to DCIS or invasive tumor cells along pseudotime.

**Extended Data Table. 1.** Assigning cell types to clusters in scFFPE-seq data.

| Cluster IDs | Assigned cell types |
| --- | --- |
| 1 | T cells |
| 2 | Stromal |
| 3 | T cells |
| 4 | T cells |
| 5 | DCIS |
| 6 | Invasive tumor |
| 7 | Macrophages |
| 8 | DCIS |
| 9 | Invasive tumor |
| 10 | Invasive tumor |
| 11 | Myoepithelial |
| 12 | B cells |
| 13 | Invasive tumor |
| 14 | Invasive tumor |
| 15 | Invasive tumor |
| 16 | Macrophages |
| 17 | Endothelial |
| 18 | B cells |
| 19 | Invasive tumor |
| 20 | Stromal |
| 21 | Macrophages |
| 22 | Plasma cells |

